# Comparing the Seasonal Diets of Buff-tailed Bumblebees and Honeybees in a Forest Landscape: A Metabarcoding Approach

**DOI:** 10.1101/2025.01.15.632979

**Authors:** Claire Gay, Précillia Cochard, Julien Thouin, Elie Morin, Fabienne Moreau, Benjamin Poirot

## Abstract

The declining diversity of pollinating insects is a major threat to ecosystem conservation, pollination services, and global food security. Honeybees (*Apis mellifera* L.) dominate managed pollination, but their dominance can affect other pollinators. Competition for resources can lead to decreased foraging success and survival rates for wild bees, especially bumblebees. This study explores the dietary composition of honeybees and buff-tailed bumblebees (*Bombus terrestris* L.) using metabarcoding techniques with three primers (ITS2, TrnLgh, and TrnLch) in Avensan, France. Primers detected different species pools, indicating a high diversity of plants visited by both species – including some false positives results inherent to metabarcoding methods. The “primer” effect was more important than the “pollinator” effect in segregating plants found. The Schoener index revealed a slight diet overlap in plant species used by honeybees and bumblebees, depending on the primer. Correspondence analyses showed a high segregation between species associated with honeybees or with bumblebees, regardless of the primer. The metabarcoding technique was found to be accurate in separating pollinator food niches, despite some biases of this technique: this result is not comparable with previous literature studying the diets of these two species, as traditional field studies are needed to complement it and overcome these biases. To conclude, this study provides a fast and inexpensive approach to study pollinators’ floral resources sharing in the same geographical area and time scale, and provide insights to improve metabarcoding effectiveness in order to better describe diet niches.

## I. Introduction

The dwindling diversity of pollinating insects, a phenomenon well-documented by numerous studies (Potts et al. 2010), poses a significant threat to ecosystems conservation, pollination service and consequently global food security (Potts et al. 2016). Several factors contribute to this decline, including the scarcity of floral resources. Indeed, plants provide essential resources for pollinators, such as pollen and nectar.

Among the most recognized pollinators is the honeybee, *Apis mellifera* Linnaeus (1758). The increasing demand for pollination services and conservation efforts has led to a proliferation of managed honeybee colonies, making them a dominant pollinator for many plants (Herrera 2020). Their dominance is attributed to their generalist foraging behavior, social organization, ease of management, high visitation rates, and efficient communication of resource locations (Crane, 1990; Rader et al., 2009; Von Frisch, 1965). However, while honeybees are valuable pollinators, they are not the sole providers of this essential ecosystem service. Other pollinators can be equally or even more efficient in certain contexts (Breeze et al. 2011).

The dominance of honeybees can negatively impact the availability of resources for other pollinating insects. The competition for floral resources between honeybees and other pollinators has become a growing concern: exploitation competition arises when one or more species deplete resources, limiting their availability for others. The overlap in ecological niches between honeybees and other pollinators can exacerbate this competition. Some recent practices in urban or agricultural landscapes, such as increasing the number of honeybee hives, can have detrimental effects on other pollinator populations by reducing their access to resources (Geslin et al. 2017, Geldmann and González-Varo 2018). Studies have shown that the presence of honeybee colonies can lead to decreased foraging success, reduced niche width, and lower survival rates for wild bees, especially bumblebees (Goulson and Sparrow, 2009; Hudewenz and Klein, 2015; Henry and Rodet, 2018; Ropars et al., 2019). Goulson & Sparrow (2009) noted that the size of workers of several species of bumblebees decreased significantly with the presence of *A. mellifera* hives nearby, with the hypothesis that individuals may receive fewer resources at the larval stage due to competition, affecting their development. Thus, in semi-natural habitats, bumblebees and solitary bees use almost half of the floral resources also used by the honeybee (Steffan-Dewenter and Tscharntke 2000). Indeed, over half of the studies examining the interaction between honeybees and other bees have reported negative effects of honeybees due to competition for resources (Mallinger et al. 2017).

But *A. mellifera* is also facing threats: Colony Collapse Disorder (CCD) played a pivotal role (vanEngelsdorp et al. 2009). CCD has profound implications for both agricultural productivity and ecosystem health. To address these challenges, strategies are being implemented to promote the populations of wild bees, including bumblebees. These efforts are essential for maintaining pollination services and ensuring food security. Studies have consistently demonstrated the superior pollination efficiency of wild bees, especially bumblebees, compared to honeybees (Javorek et al. 2002, Frier et al. 2016, Howlett et al. 2019). For instance, research by Frier et al. (2016) and Howlett et al. (2019) found that bumblebees are capable of collecting significantly more pollen per individual than honeybees. Javorek et al. (2002) and Frier et al. (2016) reported that bumblebees deposit substantially more pollen on some flowers than honeybees: honeybees need to do four times as many visits to achieve the same pollination efficiency. Similarly, bumblebees are more efficient in terms of pollen deposition and visit speed (Frier et al. 2016).

It is well known that bumblebees and honeybees have got very close diets. Thomson (2006) found a niche overlap of 80-90% between *Apis* and *Bombus* species during dearth periods in the United-States (i.e. when food resources are limited). Gay (2023) further observed that 40-55% of plant species were commonly foraged by both *A. mellifera* and *Bombus terrestris* Linnaeus (1758) in South-West France during the spring, and that this dietary overlap increased to nearly complete during the summer months. Like the honeybee before it, the generalist behaviour of the bumblebee *B. terrestris* has led humans to make it a domestic species in certain cases, notably for the pollination of tomatoes or strawberries (Velthuis and van Doorn 2006), even to the point of being established outside its natural distribution area (Goulson and Hanley 2004, Schmid-Hempel et al. 2007).

While most studies on the diet of these two species rely on traditional field methods like sweep netting (Gay et al. 2024), these approaches often demand significant expertise and time commitment. Despite their effectiveness, such methods face limitations in terms of cost, sampling effort, and statistical power, hindering their widespread adoption outside of academic research. New molecular tools, known as “metabarcoding”, have emerged to analyze the composition of pollinator diets using primers specially designed to recognize the floral species visited (Laube et al. 2010, Pornon et al. 2016). To describe the honeybee’s dietary niche – and to a lesser extent that of bumblebees –, most studies focus on pollen metabarcoding (e.g. Baksay et al. 2020; Bontšutšnaja et al. 2021; Piko et al. 2021) as a follow-up to the melissopalynological analyses used for several decades (Louveaux et al. 1978). However, more and more are looking for traces of flower DNA in other matrices such as honey (Bruni et al. 2015, Hawkins et al. 2015). This type of molecular analysis offers a more sensitive and reproducible approach than traditional microscopy for identifying plants visited by pollinators (Hawkins et al. 2015). It can detect a greater number of plant species and provide more accurate results than mellissopalinology, as it can identify not only pollen plants but also those that provide nectar (Prosser and Hebert 2017). Furthermore, pollen and nectar detected by metabarcoding in honeybees’ diets have been shown to be able to provide information about the plants growing within their flight radius (Galimberti et al. 2014, Milla et al. 2021) and could thus be used to determine the occurrence of plants of interest (Bell et al. 2016) – unfortunately, there is no data to support the same conclusion for bumblebees.

However, while increasingly obvious biases are emerging regarding the use of metabarcoding to determine pollinator diet, the lack of universal markers for all plants seems to be one of the most recurring problems (Piñol et al. 2019). When using DNA metabarcoding to study the floral composition of honey, the choice of markers is crucial. These markers must: be universal (the primers used to amplify the DNA must be designed to work with a wide spectrum of species), offer suitable discriminatory power (the region of the genome targeted by the primers must be able to differentiate between the species present) and be based on a solid reference database (the quality and completeness of the database is decisive) (Hawkins et al. 2015).This is why, in the present study, we develop our analyses using three different primers which are ITS2, TrnLgh and TrnLch. ITS2 is the ribosomal internal transcribed region 2 and TrnL is the P6 loop region of the leucine transfer RNA gene (Milla et al. 2021). ITS2 is particularly useful for distinguishing different type of plants, especially at the genus level thanks to its heightened sensitivity (Richardson et al. 2015). As for the P6 loop of TrnL, it is short and easy to amplify, making it effective for identifying plants present in samples such as pollen grains or honey, where DNA could be highly degraded (Pornon et al. 2016). In addition to the bias of the primers used and their number, there is also a bias in the assignment of plant species with metabarcoding: it is a complex method that requires attention to false positives (Cuff et al. 2022, Drake et al. 2022, Quaresma et al. 2024).

Indeed, this study proposes a novel approach to investigate the dietary composition of honeybees and bumblebees – a species usually less studied by metabarcoding –, as well as their floral resources sharing in the same geographical area and in the same time scale, through metabarcoding. By employing molecular barcoding techniques on honey and nectar samples, we sought to non-lethally characterize the plant diversity within the diet of *A. mellifera* and *B. terrestris* in their natural distribution area (De La Rúa et al. 2009, Lecocq et al. 2016), evaluate its effectiveness and biases when comparing two floral diets, and thus characterize the metabarcoding potential to infer the niche overlap between pollinators – a prevalent concern in the literature (e.g. Geslin et al., 2017; Henry and Rodet, 2018).

## II. Materials & Methods

### Sampling site

Our study site was located in the Nouvelle-Aquitaine region, southwest France, in the small town of Avensan which is characterized by a low population density (59 people per sq. km). This city has an oceanic climate, with high rainfall in autumn and winter, abundant sunshine, mild winters, and sea breezes. In the countryside of Avensan, two bumblebee hives in water-resistant cardboard (*Bombus terrestris* Linnaeus, 1758) and two honeybee wooden hives (*Apis mellifera* Linnaeus, 1758) were placed at a distance of 10 m from each other, at the end of February 2024. The location of the hives was exactly at the crossroads of coniferous forest, arable land and shrubby forest. Within a radius of 3 km (the foraging radius of honeybees), the chosen location to install these bumblebee and honeybee hives was described by: 1191.3 ha of coniferous forest (42.2%), 317.4 ha of shrub vegetation (11.2%), 288.4 ha of mixed forest (10.2%), 586.4 ha of deciduous forest (20.8%), 304.4 ha of grassland (10.8%), 126.2 ha of artificial areas such as urban areas or ground-mounted photovoltaic panels (4.5%) and 8.2 ha of roads (0.3%) (Fig. 1).

**Figure 1.**
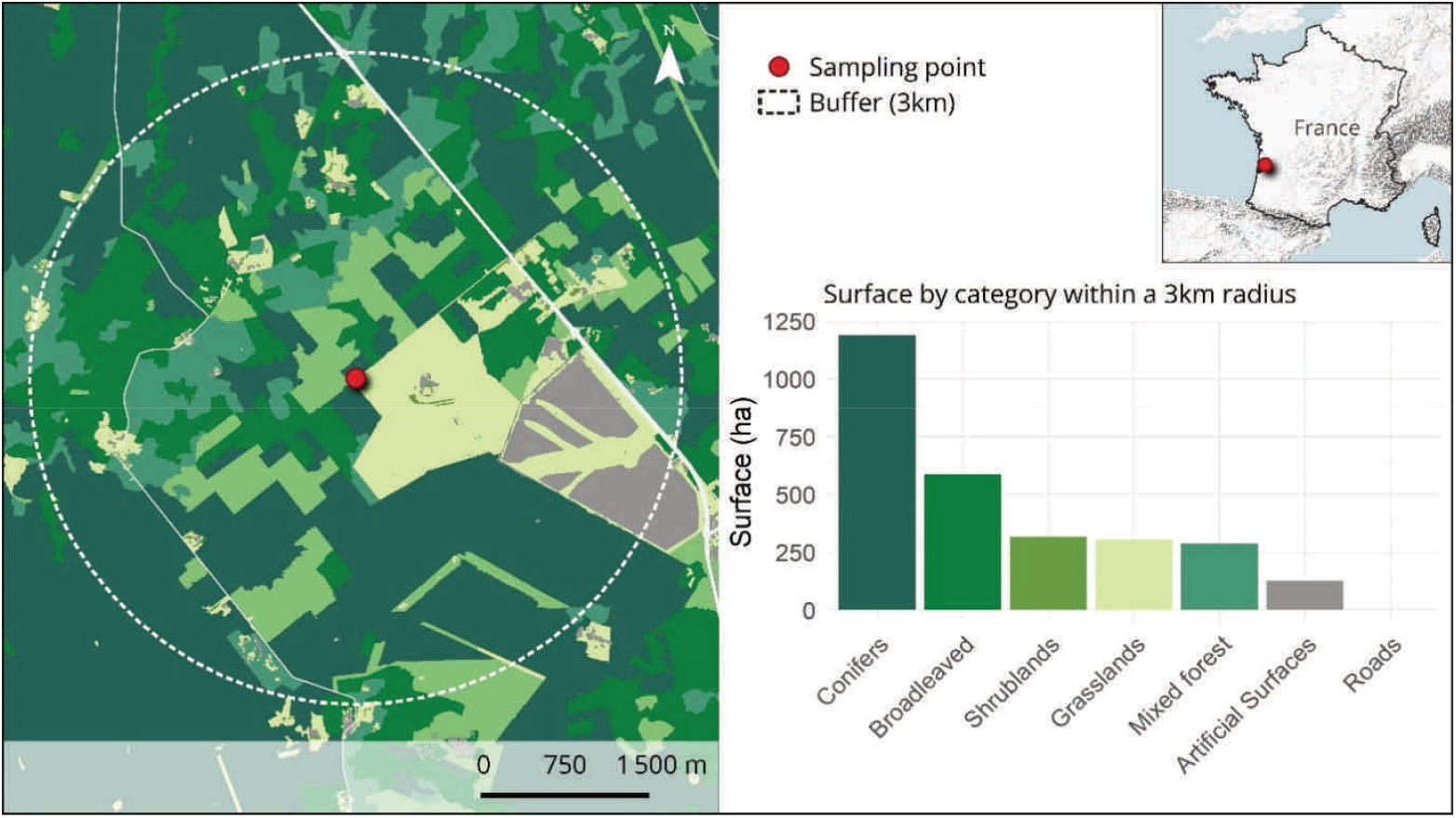
(a) Situation map of the study area, in western France (Avensan; GPS coordinates rounded to two decimal places: 44.98°N, -0.77°E) with a buffer of 3 km (b) Histogram of land use surfaces within a radius of 3 km (corresponding to the flight radius of *Apis mellifera*) using seven categories: conifers, broadleaved plants, shrublands, grasslands, mixed forest, artificial surfaces, roads.

### Sampling process and rounds

We left the hives in place for a month before any sampling, to ensure that the composition of the diet would be predominantly flowers foraged on site. In early spring, the first samples were taken. For honeybees, in May 2024, five mL of honey were collected from the frames of both hives using a sterile swab and nitrile gloves, from several frames and several locations on the frame. For bumblebees, in April 2024, ten mL of nectar mixed with pollen and cup construction residues were collected from both hives using a sterile swab and nitrile gloves (as they do not produce honey). The same sampling process was done each month - the beginning and the end of the monitoring depending on the level of activity and the peak of activity of both species according to the literature (Odoux et al. 2014).

### Metabarcoding analysis

After the samples were frozen in liquid nitrogen, ceramic beads were used on a RETSCH Mixer Mill 200 to mechanically disrupt the cells. A second cell disruption was carried out using heat treatment and a SDS based buffer. Following potassium acetate precipitation of proteins, the DNA was precipitated in isopropanol. As in the previous literature on pollen DNA (e.g. (Milla et al. 2021), PCR amplification was performed for TrnLch (i), TrnLgh (ii) and ITS2 (iii), respectively with the following primers: (i) TrnLch-F: CGAAATCGGTAGACGCTACG, TrnLch-R: CCATTGAGTCTCTGCACCTATC, (ii) TrnLgh-F: GGGCAATCCTGAGCCAA, TrnLgh-R: CCATTGAGTCTCTGCACCTATC, (iii) ITS2-F: GACTCTCGGCAACGGATATC, ITS2-R: TCCTCCGCTTATTGATATGC. The Illumina Nextera XT Index Kit v2 was used for the library preparation. Following a qPCR and fragment analyzer quality verification, the library was sequenced on a MiSeq Illumina sequencer. Internal scripts based on the FROGS v.4.1 pipeline were used to examine the raw data (Escudié et al. 2018). First, the pair-end readings were merged, and then the primers were trimmed to demultiplex them. Swarm software was used to acquire the Operational Taxonomic Units (OTUs) (Sneath and Sokal 1973) clustering with a set distance of one (Mahé et al. 2014). This caused the generation of OTUs near Amplicon Sequencing Variants (ASVs) (Eren et al. 2013). (i) The chimera was removed using filters (Haas et al. 2011), and (ii) low abundant clusters (abundance < two reads from all the samples tested: singletons, as in Oliverio et al. 2018) were eliminated to limit “false positive” identifications. The taxonomic assignment was carried out by searching the NCBI Nt database (updated in November 2023) using the blastn technique from the BLAST program v.2.13.0+ (Altschul et al. 1990).

### Statistical analysis

We estimated the completeness of DNA identification of flowers visited by bumblebees and honeybees through the three primers (ITS2, TrnLgh and TrnLch), using the Jackknife 1 estimator and Chao estimator of asymptotic species richness ((Heltshe and Forrester 1983, Chacoff et al. 2012)). We calculated the Jackknife 1 and Chao 2 estimators using the R package *vegan* and function *poolaccum* (v2.5.7; Oksanen et al., 2020) and assessed the percentage completion of the molecular flower species sampling (ratio between observed and estimated value).

We then compared the overall percentages of overlap between the floral species and the floral genera found in the bumblebee and honeybee diets using descriptive Venn diagrams using the R package *ggVennDiagram* (v1.5.2; Gao et al., 2021), which also showed the difference of results between the three primers used in the DNA analysis.

To describe seasonal trends in the number of species visited by bumblebees and honeybees, we compared the number of interaction partners (flower species) of bumblebees in April, May and June with those of honeybees in May, June, and July, by representing the mean values (with standard error, se) of the number of flowers visited regardless of the primer. The values obtained were compared using ANOVA and after checking for normality and homoscedasticity of data.

To investigate the identity of the flower species visited and its consequences, we calculated a dietary overlap index using the *spaa* package (v0.2.2, Zhang & Ma, 2014) to determine the most likely niche overlap between both pollinator species all along the studied period (Schoener 1970). This method has been used in recent studies on dietary overlap (e.g. Hilgers et al., 2018).

A correspondence analysis (CA) was done using R package *ca* (Nenadic and Greenacre 2007) for each primer separately. This approach was applied to a contingency table, between the categorical variables of pollinator type for a given sampling month and the flower species visited identified by metabarcoding. As the quantity of reads found per plant identified in the diets varied according to pollinator group, a comparison between these groups was obtained by expressing the response frequencies in relation to their respective totals (David 2017). Tabular data were represented graphically using a biplot in the form of a point cloud on two perpendicular coordinate axes (Greenacre 2007). By degrading the taxonomi resolution, we also run CA at the genus level to limit “false positive” species of plants.

All analyses were run with *R software v*.*4*.*3*.*2* (R Core Team 2024).

## III. Results

### Effectiveness of DNA analysis in flower species detection

Using Jackknife 1 estimations, we obtained a completion rate of 56.84% for honeybees diet (35 species, estimation of 61.57) and 62.96% for bumblebees diet (71 species, estimation of 112.78) (Fig. 2). Furthermore, Chao 2 estimator value was 172.29 for honeybees, and Chao 2 estimator value was 136.45 for bumblebees: we thus obtained a completion of 20.32% for honeybees plant partners and 52.03% for bumblebees plant partners using Chao 2.

**Figure 2.**
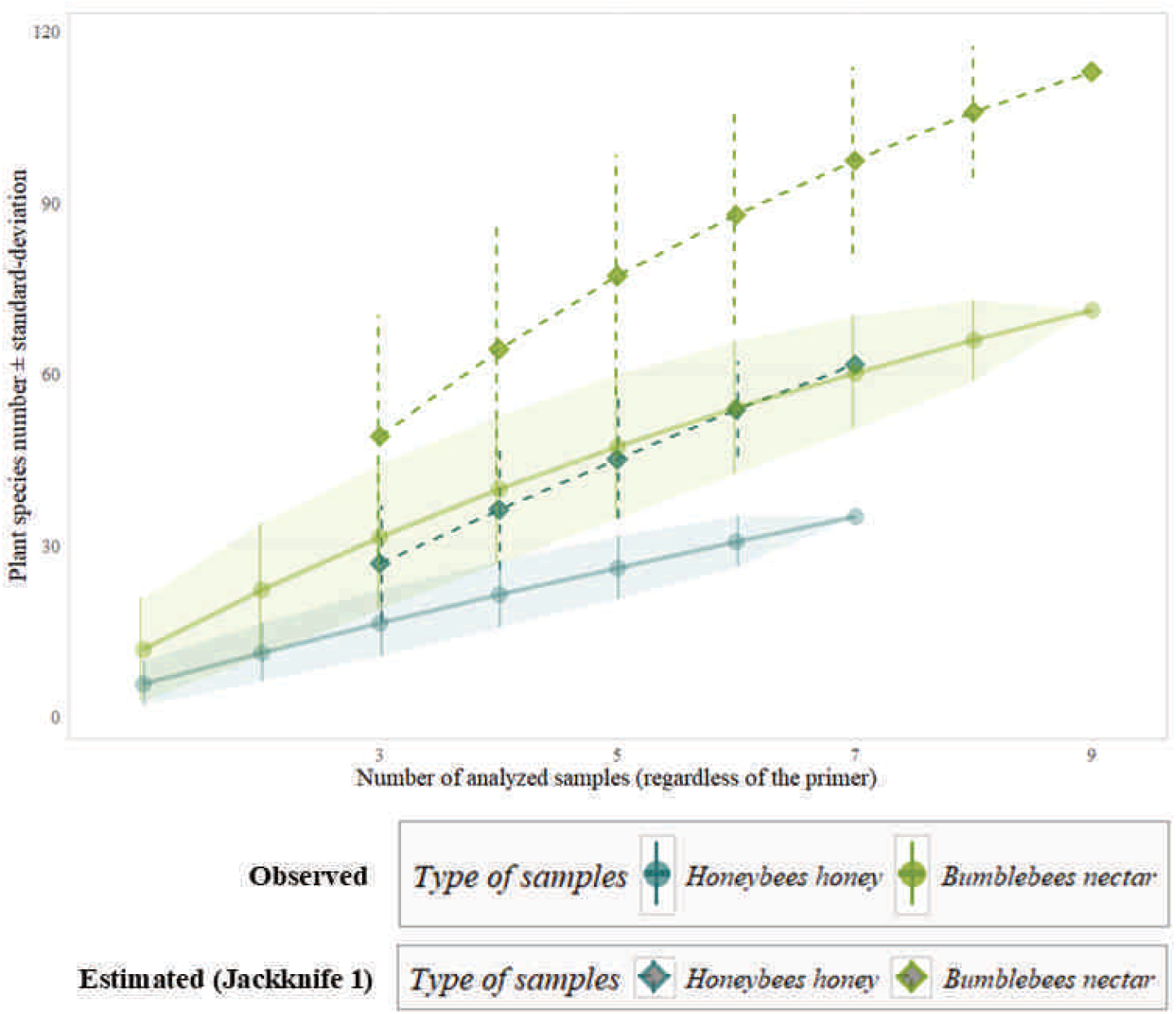
Accumulation curves of species depending on the increase of sampling units (± standard-deviation of species accumulation curves) for honeybees and bumblebees. One sampling unit is represented by one swap sample. Solid lines are observed number of species; Dashed lines are estimated number of species through Jackknife 1.

According to the Venn diagrams, the metabarcoding technique seemed adequate to segregate the diet niches of bumblebees and honeybees. Between 59.37% and 68.75% of identified plant species were only foraged by bumblebees depending on the primer (Fig. 3a), and between 15.62% and 28.13% of identified plants were only foraged by honeybees depending on the primer. Moreover, results show that between 10% and 15% of plants found in the matrices were visited by both types of pollinators (10.81% to 15.63%). The best primer to highlight the shared plants between honeybees and bumblebees was TrnLch (five species) but it was ITS2 to highlight the plants specific to the bumblebee (23 species) and the plants specific to the honeybee diet (10 species). At the genus level, repartition of floral diets was approximately the same as for the species level: between 59.37% and 66.66% of identified plant species were only foraged by bumblebees depending on the primer and between 10.71% and 28.13% of identified plants were only foraged by honeybees depending on the primer (Fig. 3b).

**Figure 3.**
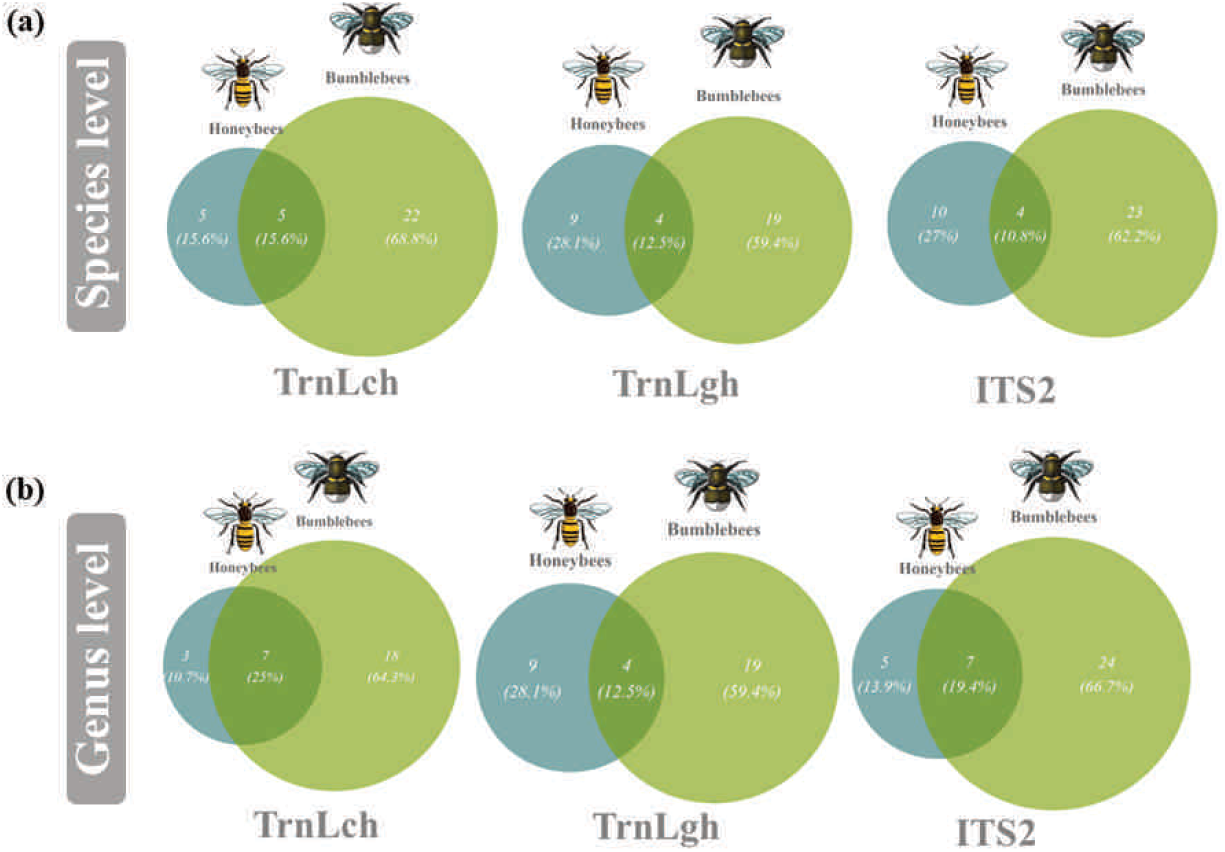
Venn diagram showing the overlapped detected (a) plant species and (b) plant genus and the respective percentages they represent between honey samples for honeybees and nectar samples for bumblebees; with TrnLch DNA primer (left), with TrnLgh DNA primer (middle) and with ITS2 DNA primer (right).

### Seasonal variation in number of visited plants by honeybees and bumblebees

Seasonal trend in number of species detected in honeybees and bumblebees diet (with the DNA primer as a source of variability) was represented in Fig.4. In May, the number of species visited by honeybees was 8.667 ± 2.667 (mean ± standard-error), while it was 5.333 ± 1.856 in April for bumblebees and 23.667 ± 2.404 in May for bumblebees. In early spring, in May, the difference in number of plants species detected in the diet of honeybees and in the diet of bumblebees was significant (F_1,6_ = 23.520, p-value = 0.003). During the summer, the average number of species found for honeybees was 1.500 ± 0.500 in June, 5.500 ± 0.500 in July, and 6.000 ± 2.517 for bumblebees in June. In June, we did not found any significant difference in the number of species visited per pollinator type (F_1,4_ = 3.379, p-value = 0.140).

**Figure 4.**
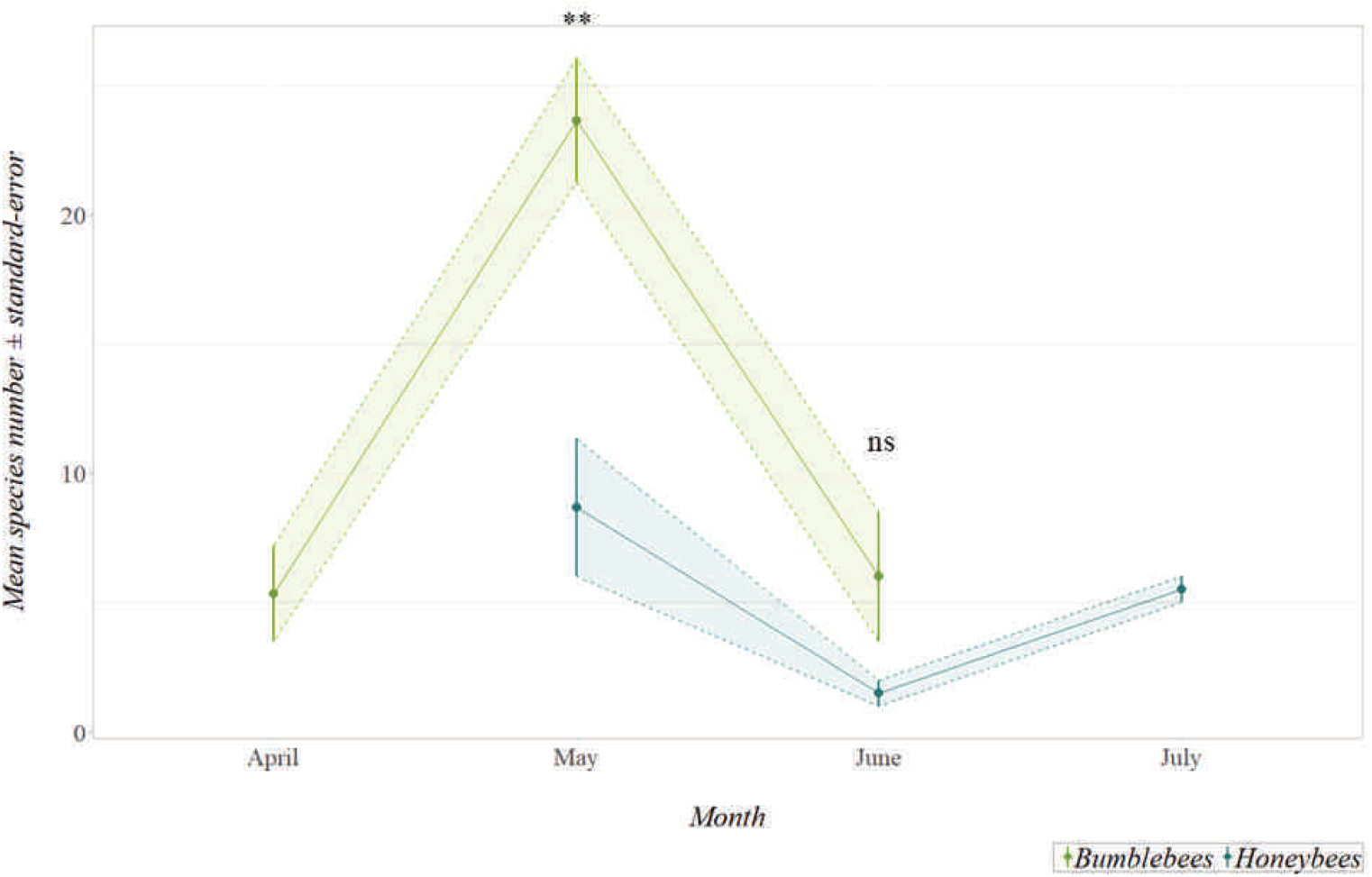
Number of visited plant species (number of different plant species detected in honey for honeybees and nectar for bumblebees) throughout the spring (April, May, June, July). Data are presented as mean ± se (standard error). Analyses of variances were run on each month to compare both insect diets. If p-value < 0.001 then ***; If 0.001 $p-value < 0.01 then **; If 0.01 $p-value < 0.05 then *; If 0.05 $p-value < 0.1 then ‘.’ Green points and line: Bumblebees; Turquoise points and line: Honeybees.

### Differences in floral pool composition in honeybees and bumblebees diets

After studying the difference in species composition between the different primers (ITS2, TrnLgh and TrnLch), we chose to do the species identity analysis by separating primers from each other. Indeed, the primers have detected pools of different species during metabarcoding, whether in bumblebees or honeybees. ITS2 detected 21 species alone (i.e. that were not detected by the other primers) in the diet of bumblebees and 10 in that of honeybees. TrnLgh detected 14 species of flowers in bumblebees and eight species in honeybees that were not detected by the other primers. Finally, TrnLch detected 18 species alone in bumblebees and three species in honeybees. The “primer” effect was more important than the “pollinator” effect in segregating the plants found. For instance, there were more species found in common between bumblebees *via* TrnLch and honeybees *via* TrnLch (five species) than species in common between bumblebees *via* TrnLch and *via* TrnLgh (three species). As well, there were more species found in common between bumblebees *via* ITS2 and honeybees *via* ITS2 (four species) than species in common between bumblebees *via* ITS2 and *via* TrnLgh or TrnLch (respectively two and zero species). Note that there were no flower species detected in common by the three primers, either in bumblebees or honeybees **(see ESM1 in Appendix A)**.

The Schoener index revealed a very slight diet overlap in plant species used by honeybees and bumblebees, depending on the primer (S_index_max=0.221 for ITS2, S_index_min=0.026 for TrnLgh) (Tab.1). It did not increase much for ITS2 when focusing on shared plant genus (S_index_=0.229), even though the diet overlap was higher between honeybees and bumblebees when using TrnLch primer (S_index_=0.361 for TrnLch, S_index_=0.122 for TrnLgh). The diet overlap between honeybees and bumblebees was in comparison quite high (S_index_=0.368 for ITS2, S_index_=0.407 for TrnLch, S_index_=0.323 for TrnLgh) when focusing on shared plant families.

The correspondence analysis showed different results for each primer. With the ITS2 primer (component 1: 89.621%; component 2: 10.080%), *Pinus contorta* [American but plausible species due to importation on Atlantic coast in the 1950s; **see ESM2 in Appendix A for all species presence plausibility**] (40.540%), *Lamium purpureum* (16.212%) and *Trifolium incarnatum* (16.331) were the three best contributors to the inertia of component 1, whereas they were *Ulex europaeus* (58.127%), *Lamium purpureum* (0.001%) and *Plantago ovata* (10.804%) for component 2 (Fig.5.a.). With the TrnLgh primer (component 1: 26.067%; component 2: 23.800%), *Galega officinalis* (50.001%), *Calluna vulgaris* (12.797%) and *Crepis sancta* (11.171%) were the three best contributors to the inertia of component 1, whereas they were *Rhamnus crenata* [implausible species – degraded to genus level: *Rhamnus sp*.] (73.558%), *Hypericum androsaemum* (14.545%) and *Crepis sancta* (5.746%) for component 2 (Fig.5.b.). With the TrnLch primer (component 1: 41.888%; component 2: 29.786%), *Ranunculus macranthus* [implausible species – degraded to genus level: *Ranunculus sp*.] (68.252%), *Lonicera fragrantissima* (16.653%) and *Salvia rosmarinus* (3.666%) were the three best contributors to the inertia of component 1, whereas they were *Persea schiedeana* [implausible species or imported – degraded to genus level: *Persea sp*.] (32.729%), *Ophrys insectifera* (8.032%) and *Erica arborea* (7.033%) for component 2 (Fig.5.c.). Correspondence analyses also showed a high segregation between species associated with honeybees and species associated with bumblebees, regardless of the primer. Indeed, the further represented species from the origin and thus the most discriminated were those associated with honeybees, which was usually strongly associated with a small number of taxa: for instance, on component 1, honeybees cos2 was 0.999 in May for ITS2, 0.999 in May for TrnLgh or 0.979 in May for TrnLch. On the contrary, bumblebees nectar samples were weaker represented on components: on component 1, best bumblebees cos2 was 0.979 in May for ITS2 (but 0.080 in April and 0.018 in June), 0.270 in June for TrnLgh or 0.388 in June for TrnLch. Moreover, bumblebees diet was quite similar except when identifying plants with TrnLch primer which better segregate their diets. Their diets were always associated with a huge number of plants on both components, as for honeybees with ITS2 (plants’ DNA detected only for July – no results for other months). With ITS2, honeybees diet was strongly associated with several *Pinus* species as well as *Trifolium, Quercus, Ilex* and *Prunus* species in May. Nevertheless, it was not as diverse for honeybees diets with TrnLgh and TrnLch. With TrnLgh, honeybees diet was strongly associated with several *Calluna* or *Chelidonium* species in May, with *Rhamnus* species in June, and with *Cornus* and *Mercurialis* species in July. With TrnLch, honeybees diet was strongly associated with *Ranunculus macranthus, Genista tridentata* and *Medicago marina* in May, with *Cucumis melo* and *Lonicera fragrantissima* in June, and with *Tripleurospermum maritimum* and *Cornus florida* [an imported species] in July. Correspondence analyses at degraded taxonomic level (i.e. genus, to limit impact of “false positive” species) are available in **ESM3 in Appendix A**.

**Table 1.** Schoener index values (dietary niche overlap) according to the insect taxonomic group and the DNA primer used for the analyses. Values between zero (no dietary niche overlap) to one (total dietary niche overlap).

| Taxonomic level |  | Primer |  |  |
| --- | --- | --- | --- | --- |
|  |  | ITS2 | TrnLgh | TrnLch |
|  | Species | 0.221 | 0.026 | 0.034 |
|  | Genus | 0.229 | 0.122 | 0.361 |
|  | Family | 0.368 | 0.323 | 0.407 |

**Figure 5.**
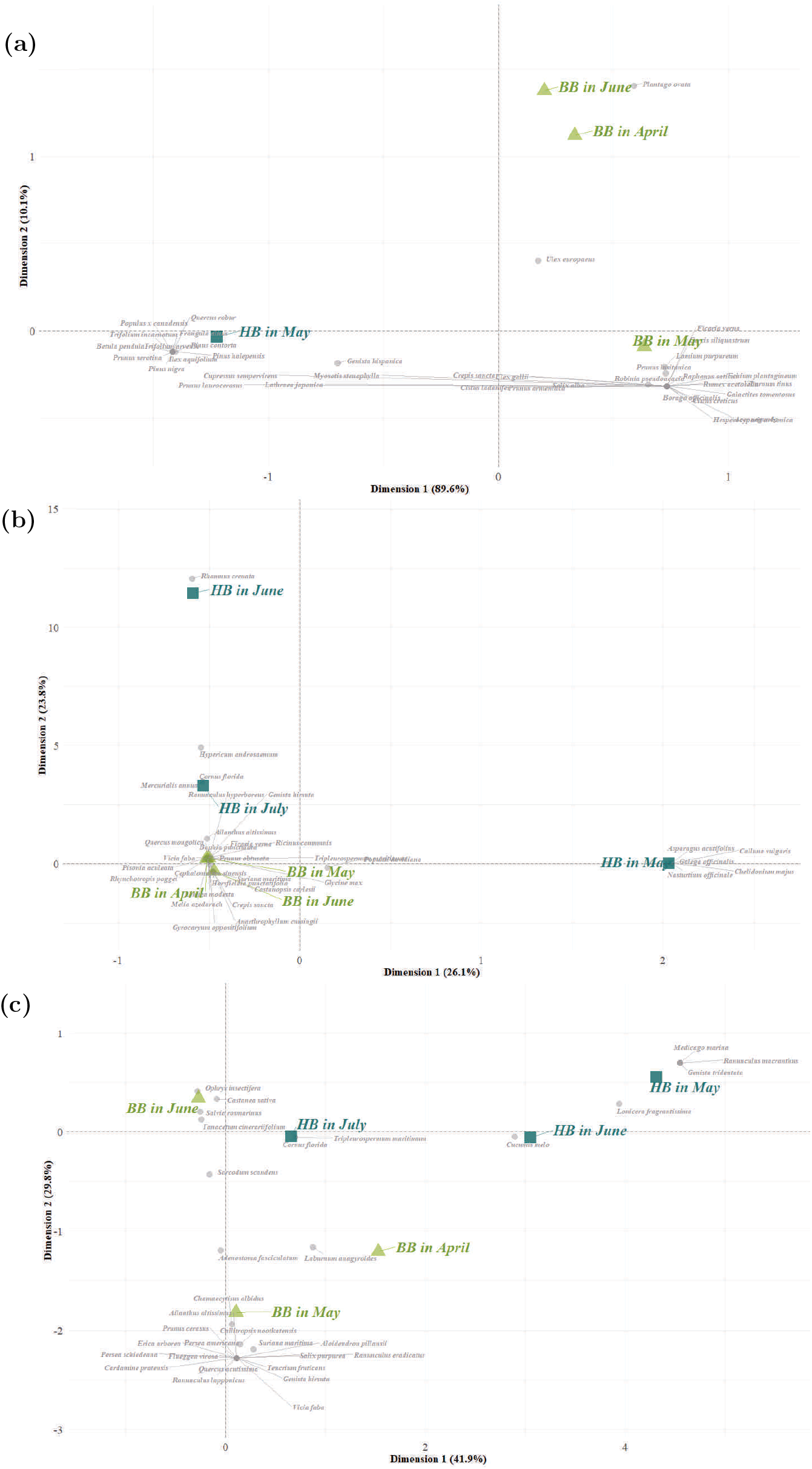
Biplot of the correspondence analysis for (a) ITS2 DNA primer, (b) TrnLgh DNA primer and (c) TrnLch DNA primer. Green triangles: Bumblebees nectar samples in April, May or June; Turquoise squares: Honeybees honey samples in May and July; Grey points: Plant species detected by metabarcoding analyses. 10

## IV. Discussion

The detection of a high number of taxa in the plants visited by bumblebees and honeybees indicates that our study site is probably characterized by a high diversity of floral species despite the ubiquitous planted pine forests on the Atlantic coast, which is known to have some negative effects on biodiversity (Carnus et al. 2006) especially in our study area (Jouveau et al. 2022). This potential diversity of floral species could be explained by a heterogeneous landscape – adjacent to the pine forest – showing agricultural fields, deciduous forests as well as shrublands. We emphasized the presence of emblematic species from the study region landscape such as the gorse *Ulex europaeus*, the heather *Calluna vulgaris* or the locally protected species *Medicago marina* already known to be present in the Médoc and Entre-Deux-Mers locations (Favennec 2002, Ministère de l’aménagement du territoire et de l’environnement 2002). However, we did not detect all the species visited at all according to the estimates of Jackknife 1 and Chao 2 – but sampling diets of pollinators is always difficult and other studies tend to detect less than three quarters of the plants visited (Chacoff et al. 2012) or need a long-term sampling effort to reach high completion rates (Gay et al., 2024). On the one hand, the sampling effort should be increased, as the plateau of the rarefaction curve has not been reached. On the other hand, the number of species detected may have been underestimated due to the low efficiency of the primers used for detecting certain plant species (Hawkins et al. 2015) or, on the contrary, overestimated by the generation of “false positive” species, which has repercussions on the calculation of Jackknife 1 and Chao 2 values – themselves overestimated (Drake et al. 2022). Indeed, describing the ecological niche in its entirety or in its veracity (i.e. to detect all the plants in the diet with certainty) could then substantially change our conclusions. It has already been shown that metabarcoding can detect floral taxa that are abundant in honey, but not less abundant plants (Hawkins et al. 2015), potentially explaining our poor completion of diet composition – in addition to a small number of samples. Nevertheless, metabarcoding makes it possible to identify plants with which the pollinators studied have actually been in contact by leaving traces of DNA, which is not possible with major part of non-lethal methods (such as botanical studies). To go further, melissopalynology (Louveaux et al. 1978) could provide similar or even more accurate results than DNA analyses, but the economic cost and human resources required for melissopalynology are an argument in favor of metabarcoding methods (Bell et al. 2016, Milla et al. 2021).

Using three different DNA primers, we obtained similar results in terms of number of visited floral species but not in terms of species identity. The three primers, although designed to detect plant DNA, do not have the same taxonomic resolution as a result of variable base pair length. The results of DNA analyses vary greatly depending on the primer used, highlighting the need to choose appropriate primers (Prosser and Hebert 2017), and the benefits of using and comparing different primers. But some progress is expected in future literature dealing with pollinators diet through metabarcoding: we have encountered errors in the identification of species by this method (probably due to lacks of knowledge or errors in DNA databases, Quaresma et al., 2024), indicating species from distant geographical areas that are implausible for import (e.g. *Rhamnus crenata, Persea schiedeana*). This is a well-known drawback of metabarcoding applied to plant-pollinator interactions, which has already been documented in the literature (Cuff et al. 2022). For better species matching, we recommend only retaining species identifications with a BLAST identity score greater than 98% as applied by several other studies (e.g. Alexander et al., 2023; Webster et al., 2020) – but which we prefer not to apply in this study in order to obtain the most exhaustive possible plant composition of the diets and highlight false positive concern. It’s true that we may miss species or new identifications because of this 98% threshold, but it enables future authors to obtain a more certain list of flower species, the benefit-risk balance being in favor of applying a threshold.

Moreover, on-site botanical expertise is still necessary for studies aimed at determining the presence of a new species interaction in a given geographical area, and to establish robust pollination interaction networks. Indeed, even though we applied a sorting procedure eliminating all plant species occurrences below two DNA reads, the choice of this threshold is critical in generating artefacts (i.e. false positives), as demonstrated by Drake et al. (2022). Increasing this threshold would allow future studies to limit this bias and obtain results more consistent with literature using traditional field sampling to measure niche overlaps between pollinators – as they highlight huge niche overlaps between honeybees and bumblebees (Thomson 2006, Gay 2023).

Despite these drawbacks, the metabarcoding technique seems accurate to separate different pollinator food niches, always bearing in mind that we detected few species in the diets according to our completion rates: we did not found the same pool of foraged species between bumblebees and honeybees, which underlines the ability of DNA analyses to detect different kind of floral species. This is a new insight for metabarcoding, since only few publications have focused on several types of pollinating insects in a same study (e.g. Casanelles-Abella et al., 2022). Here, focusing on the diets of honeybees and bumblebees is an interesting aspect, given the issues raised in recent years about the competition pressure from honeybees on other pollinators (Mallinger et al. 2017). On the contrary to previous literature, we did not found a huge overlap in the diet of these two insects. While Thomson (2006) found a niche overlap of 80-90% between *A. mellifera* and *B. terrestris* and while Gay (2023) found a niche overlap of 40-55% to nearly complete according to the season between *A. mellifera* and *B. terrestris*, we did not found high niche overlap using Schoener index (Schoener 1970) – not exceeding the threshold of 0.6 emphasized in numerous studies to describe a significant overlap (Wallace and Ramsey 1983). It should be noted that our results were obtained in a forest landscape, whereas other studies often focused on open-field areas. Our study area exhibited a high diversity of semi-natural habitats, leading to a greater variety of plant species. In fact, this diversification of plants could explain the low niche overlap, as limited competition for resource access occurred between honeybees and bumblebees. This also highlights the fact that the more diverse and complex a landscape is, the more resilient and rich plant communities are. However, a potential risk of dietary niche overlap is highlighted for June on the plants detected through environmental DNA in the present study, as the number of species visited by honeybees and bumblebees has collapsed during this month. Indeed, June is the period during which the needs of honeybees hives are at their highest because the number of individuals is at its maximum within the colonies (Odoux et al. 2014). Nevertheless, it is interesting to note that the weather in this area was not clement in May, when we found significantly less floral species visited by honeybees than visited by bumblebees, with a lot of rain and cool temperatures (data from the weather station installed on the adjacent Arsac photovoltaic site), plausibly coinciding with a low number of honeybees leaving their hives to forage previous **(see ESM4 in Appendix A)**. Indeed, the foraging activity of bumblebees *B. terrestris* is less perturbed by inclement weather conditions than that of honeybees *A. mellifera* (Dag et al. 2006) and honeybees are very sensitive to changes in weather conditions within a day (Karbassioon et al. 2023). This is another argument that could explain the very low completion rate of the honeybee diet in our data.

We also observed a more generalist behavior in bumblebees than in honeybees, as they visited a wider range of floral resources in the present study. Bumblebees and honeybees are the most frequent visitors of flowers in the study area (South-West France, Gay et al., 2024), but honeybees are usually considered as those with the more generalist behavior (Giannini et al. 2015, Geslin et al. 2017). Nevertheless, it is also known that bumblebees that we used in this study, *B. terrestris*, are able of substantial foraging skills (Dafni and Shmida 1996): it is a generalist species, foraging at low temperatures, visiting deep flowers, and demonstrating buzz pollination and nectar robbing techniques (Geslin et al. 2017). They are able of collecting more pollen than honeybees in terms of individual pollination, and are more efficient in terms of pollen deposition and visit speed (Frier et al. 2016, Howlett et al. 2019).

By exploring the diets of honeybees and bumblebees – a less studied group of species – through metabarcoding, the present study provides a good but perfectible approach to study their floral resources sharing in the same geographical area as well as in the same time scale. From our results, it appears that a greater sampling effort associated with a DNA primers combination and a special attention to the sorting thresholds enables future studies to increase knowledge on the use of metabarcoding methods to compare multiple pollinators dietary niches. This could perhaps later complement botanical inventories by determining the plausible insect-pollinated plant community of a given area. Further research needs to be conducted to improve plants DNA databases, and to find a better combination of primers to cover the full and true diversity of plants pollinated by these two hymenopteran species. Finally, our results provide new insights into resource competition between honeybees and bumblebees. The low level of niche overlap observed should be further validated using metabarcoding in other environments, such as open-fields, forests, and urban areas, as plant diversity can vary significantly across these landscapes. This study also highlights the importance of increasing habitat complexity, particularly in open-field areas that have been homogenized, by incorporating agroecological infrastructures (e.g. hedgerows).

Such measures are expected to promote more stable population dynamics among pollinators and sustain robust ecosystem services essential for human activities.

## Author contributions

Conceptualization, C.G., P.C. and B.P.; Methodology, C.G., P.C., B.P., J.T. and F.M.; Formal Analysis, C.G.; Writing-Original Draft Preparation, C.G., B.P, J.T., E.M. and F.M. Writing-Review and Editing, C.G., E.M. and B.P. All authors have read and agreed to the published version of the manuscript.

## Acknowledgments

We would like to thank the teams of Engie Green^®^ Company - Agence Sud-Ouest (in particular Antoine Pouey) for funding the *Apis mellifera* honey sampling and helping to find suitable locations for the bumblebee and honeybee hives. We would also like to thank the Apilab^®^ team for funding the *Bombus terrestris* nectar sampling and their beekeepers for their help in providing equipment for monitoring the hives. Our sincere thanks to Eliot Poirot for the drawings of bees and bumblebees in Figure 3.

## Data availability

Raw amplicon reads and genome assemblies have been deposited in the *European Nucleotide Archive* under the accession number PRJEB83440.

## Conflict of interest

The authors declare that they have no conflict of interest.

## Appendix A Supporting information

Supplementary data associated with this article can be found in the online version at […].

## Notes

### Competing Interest Statement

The authors have declared no competing interest.

### Summary of Updates

Figures revised - changes in decimals in some sections of Results - new insights in discussion

