## Supplementary Materials for "Comparing the Seasonal Diets of Buff-tailed Bumblebees and Honeybees in a Forest Landscape: A Metabarcoding Approach"

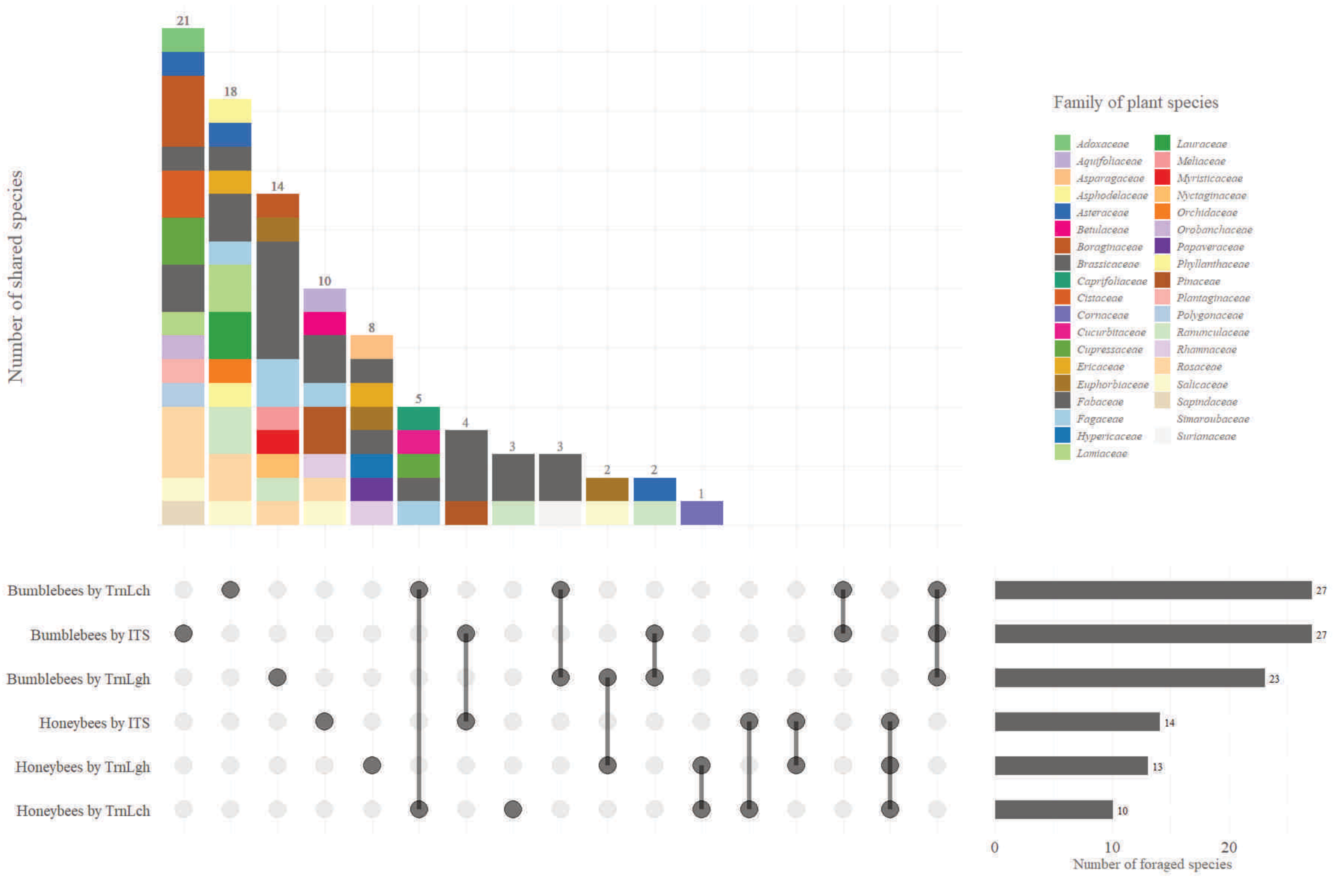


**Supplementary material 1**

Number of shared species between the different types of insects and primers. Black points represent the insect type and the primer considered in the calculation, whereas light grey points represent the insect type and the primer that are excluded from the calculation. Black bars represent the species set size of the samples. Number of shared species and the families they belong are represented by the histogram bars, above the insect and primer assemblage they represent. Insect and primer assemblages are ordered from the ones which contain most shared species, to the ones which do not contain any shared species.

### Supplementary material 2

Presence plausibility (four modalities, from gray to brown) of species extracted from environmental DNA on the Avensan site. Plausibility was checked using free data aggregation platforms and maps such as *gbif.org* and *plantnet.org*'s 'Explore' portal.

| Species | Plausibility | Note on the species |
|---|---|---|
| ***Acer negundo*** | Highly plausible | Invasive, common |
| ***Adenostoma fasciculatum*** | Highly improbable | Not introduced, Californian |
| ***Ailanthus altissimus*** | Highly plausible | Frequent invasive |
| ***Aloidendron pillansii*** | Highly improbable | South African |
| ***Anarthrophyllum cumingii*** | Highly improbable | Non-introduced, Andian |
| ***Asparagus acutifolius*** | Highly plausible | Present in the south-west of France |
| ***Baphia punctulata*** | Highly improbable | Non-introduced, tropical, African |
| ***Betula pendula*** | Highly plausible | Common |
| ***Borago officinalis*** | Possible presence | Cultivated, sometimes spontaneous |
| ***Callitropsis nootkatensis*** | Unlikely to occur | Rare ornamental, North American |
| ***Calluna vulgaris*** | Highly plausible | Common on the Atlantic coast |
| ***Cardamine pratensis*** | Highly plausible | Common |
| ***Castanea sativa*** | Highly plausible | Indigenous, common in Aquitaine |
| ***Castanopsis carlesii*** | Highly improbable | Non-introduced, Asian |
| ***Cephalomappa sinensis*** | Highly improbable | Non-introduced, Asian |
| ***Cercis siliquastrum*** | Possible presence | Ornamental |
| ***Chamaecytisus albidus*** | Unlikely to occur | Mediterranean |
| ***Chelidonium majus*** | Highly plausible | Common |
| ***Cistus creticus*** | Highly plausible | Present in the south-west of France |
| ***Cistus ladanifer*** | Highly plausible | Present in the south-west of France |
| ***Cornus florida*** | Possible presence | Ornamental |
| ***Crepis sancta*** | Highly plausible | Present in the south of France |
| ***Cucumis melo*** | Possible presence | Cultivated, sometimes spontaneous |
| ***Cupressus sempervirens*** | Possible presence | Ornamental |
| ***Echium plantagineum*** | Highly plausible | Invasive, South European |
| ***Erica arborea*** | Highly plausible | Locally present |
| ***Erica cinerea*** | Highly plausible | Indigenous |
| ***Ficaria verna*** | Highly plausible | Common in wetlands |
| ***Fitchia speciosa*** | Highly improbable | Native to the Pacific, not introduced |
| ***Flueggea virosa*** | Highly improbable | Non-introduced, pantropical |
| ***Frangula alnus*** | Highly plausible | Native to wetlands |
| ***Galactites tomentosus*** | Highly plausible | Present in the south-west of France |
| ***Galega officinalis*** | Possible presence | Introduced and sometimes spontaneous |
| ***Genista hirsuta*** | Highly plausible | Present in the south-west of France |
| ***Genista hispanica*** | Highly plausible | Very present in the region |
| ***Genista tridentata*** | Highly plausible | Present in the south-west of France |
| ***Glycine max*** | Possible presence | Cultivated |
| ***Gyrocaryum oppositifolium*** | Highly improbable | Unknown in Europe |
| ***Hesperocyparis arizonica*** | Unlikely to occur | Rare ornamental, North American |
| ***Horsfieldia punctatifolia*** | Highly improbable | Non-introduced, tropical |
| ***Hypericum androsaemum*** | Highly plausible | Common |
| ***Hypochaeris radicata*** | Highly plausible | Common |
| ***Ilex aquifolium*** | Highly plausible | Native, common |
| ***Laburnum anagyroides*** | Possible presence | Ornamental |
| ***Lamium purpureum*** | Highly plausible | Common |
| ***Lathraea japonica*** | Highly improbable | Non-introduced parasitic plant, Asian |
| ***Lennea modesta*** | Highly improbable | Non-introduced exotic |
| ***Lonicera fragrantissima*** | Possible presence | Rare ornamental |
| ***Medicago marina*** | Possible presence | Coastal dune leguminous |
| ***Melia azedarach*** | Unlikely to occur | Rare ornamental, Subtropical |
| ***Mercurialis annua*** | Highly plausible | Very common weed |

| Species | Plausibility | Note on the species |
|---|---|---|
| ***Myosotis stenophylla*** | Unlikely to occur | Uncommon |
| ***Nasturtium officinale*** | Highly plausible | Common |
| ***Ophrys insectifera*** | Highly plausible | Native |
| ***Persea americana*** | Unlikely to occur | Rare outdoors |
| ***Persea schiedeana*** | Highly improbable | Tropical |
| ***Pinus contorta*** | Possible presence | Introduced in forest plantations |
| ***Pinus halepensis*** | Possible presence | Introduced, locally present |
| ***Pinus nigra*** | Highly plausible | Often planted |
| ***Pisonia aculeata*** | Highly improbable | Not introduced, tropical |
| ***Plantago ovata*** | Possible presence | Mediterranean, rare in this area but already recorded |
| ***Populus davidiana*** | Highly improbable | Non-introduced, Asian |
| ***Populus x canadensis*** | Highly plausible | Hybrid widely cultivated in France |
| ***Prunus armeniaca*** | Possible presence | Locally cultivated |
| ***Prunus cerasus*** | Possible presence | Cultivated |
| ***Prunus grayana*** | Highly improbable | Non-introduced, Asian |
| ***Prunus laurocerasus*** | Highly plausible | Ornamental, often naturalized |
| ***Prunus lusitanica*** | Possible presence | Ornamental |
| ***Prunus obtusata*** | Highly improbable | Non-introduced, Asian |
| ***Prunus serotina*** | Highly plausible | Well-established invasive |
| ***Quercus acutissima*** | Unlikely to occur | Rarely planted, Asian |
| ***Quercus mongolica*** | Highly improbable | Rare, Asian |
| ***Quercus robur*** | Highly plausible | Native, common |
| ***Ranunculus eradicatus*** | Unlikely to occur | No local reports |
| ***Ranunculus hyperboreus*** | Highly improbable | Not adapted to the climate, Arctic |
| ***Ranunculus lapponicus*** | Highly improbable | Not adapted to the climate, Arctic/boreal |
| ***Ranunculus macranthus*** | Unlikely to occur | Non-native, rare and not cultivated locally |
| ***Raphanus sativus*** | Possible presence | Cultivated |
| ***Rhamnus crenata*** | Highly improbable | Little documented, not introduced |
| ***Rhynchotropis poggei*** | Highly improbable | Non-introduced, tropical, African |
| ***Ricinus communis*** | Possible presence | Sometimes present in the south-west of France |
| ***Robinia pseudoacacia*** | Highly plausible | Naturalized invasive, widespread |
| ***Rumex acetosella*** | Highly plausible | Common |
| ***Salix alba*** | Highly plausible | Native to wetlands |
| ***Salix purpurea*** | Highly plausible | Native |
| ***Salvia rosmarinus*** | Possible presence | Cultivated, sometimes spontaneous |
| ***Sarcodum scandens*** | Highly improbable | Non-introduced, tropical |
| ***Suriana maritima*** | Highly improbable | Not introduced, tropical |
| ***Tanacetum cinerariifolium*** | Possible presence | Cultivated on a small scale |
| ***Teucrium fruticans*** | Possible presence | Ornamental, Mediterranean |
| ***Trifolium arvense*** | Highly plausible | Common |
| ***Trifolium incarnatum*** | Highly plausible | Cultivated, frequently observed |
| ***Trifolium repens*** | Highly plausible | Extremely common |
| ***Tripleurospermum maritimum*** | Highly plausible | Common on the Atlantic coast |
| ***Ulex europaeus*** | Highly plausible | Native, very common |
| ***Ulex gallii*** | Highly plausible | Common |
| ***Viburnum tinus*** | Possible presence | Frequent ornamental, Mediterranean |
| ***Vicia faba*** | Possible presence | Cultivated |

### Supplementary material 3

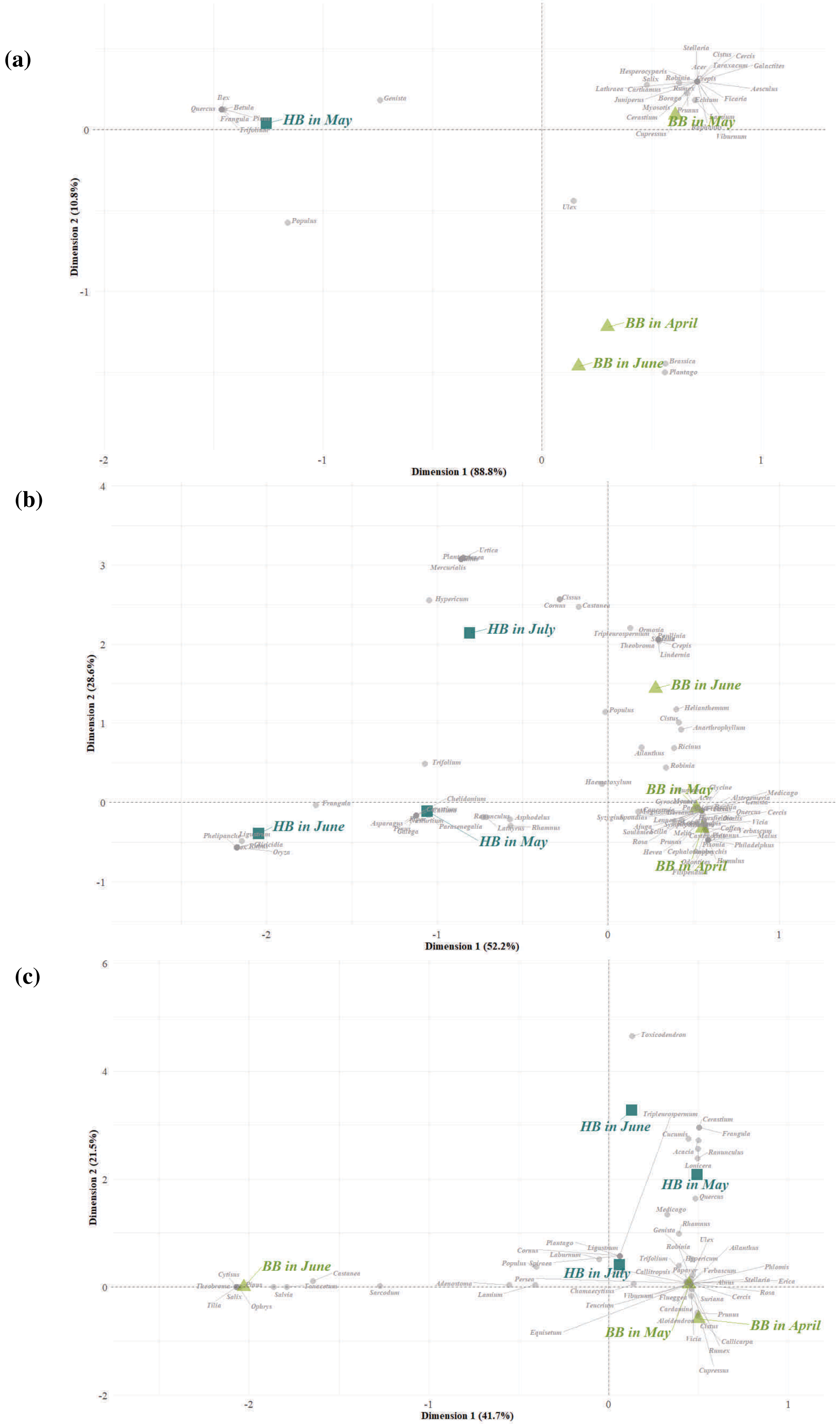


Biplot of the correspondence analysis for (a) ITS2 DNA primer, (b) TrnLgh DNA primer and (c) TrnLch DNA primer. Green triangles: Bumblebees nectar samples in April, May or June; Turquoise squares: Honeybees honey samples in May, June and July; Grey points: Plant genus detected by metabarcoding analyses.

**Supplementary material 4**

Meteorological data of local station from March to September. Device installed at the Arsac photovoltaic site, monitored by the Engie Green® Company team and adjacent to the study site. Shaded lines indicate months when sampling did not take place.

| **Month** | **Weather conditions** | |
|---|---|---|
| | *Monthly total rainfall (mm)* | *Monthly average temperature (°C)* |
| *March* | 196.9 | 10.55 |
| *April* | 68.4 | 16.85 |
| *May* | 191.8 | 14.77 |
| *June* | 103.2 | 19.12 |
| *July* | 16.2 | 21.45 |
| *August* | 14.5 | 21.61 |
| *September* | 16.5 | 16.33 |
